# Test Gene-Environment Interactions for Multiple Traits in Sequencing Association Studies

**DOI:** 10.1101/710574

**Authors:** Jianjun Zhang, Qiuying Sha, Han Hao, Shuanglin Zhang, Xiaoyi Raymond Gao, Xuexia Wang

## Abstract

The risk of many complex diseases is determined by a complex interplay of genetic and environmental factors. Data on multiple traits is often collected for many complex diseases in order to obtain a better understanding of the diseases. Examination of gene-environment interactions (GxEs) for multiple traits can yield valuable insights about the etiology of the disease and increase power in detecting disease associated genes. Most existing methods focus on testing gene-environment interaction (GxE) for a single trait. In this study, we develop novel approaches to test GxEs for multiple traits in sequencing association studies. We first perform transformation of multiple traits by using either principle component analysis or standardization analysis. Then, we detect the effect of GxE for each transferred phenotypic trait using novel proposed tests: testing the effect of an optimallyweighted combination of GxE (TOW-GE) and/or variable weight TOW-GE (VW-TOW-GE). Finally, we employ the Fisher’s combination test to combine the p-values of TOW-GE and/or VW-TOW-GE. Extensive simulation studies based on the Genetic Analysis Workshop 17 data show that the type I error rates of the proposed methods are well controlled. Compared to the existing interaction sequence kernel association test (ISKAT), TOW-GE is more powerful when there are only rare risk and protective variants; VW-TOW-GE is more powerful when there are both rare and common risk and protective variants. Both TOW-GE and VW-TOW-GE are robust to directions of effects of causal GxEs. Application to the COPDGene Study demonstrates that our proposed methods are very powerful.

## Introduction

The risk of many complex diseases is determined by a complex interplay of genetic and environmental factors. For example, anthracyclines are one of the most effective classes of chemotherapeutic agents currently available for cancer treatment. The therapeutic potential of anthracyclines, however, is limited because of their strong dose-dependent relation with progressive and irreversible cardiomyopathy leading to congestive heart failure. Both gene hyaluronan synthase 3 (HAS3) and gene CUGBP Elav-like family member 4 (CELF4) exert modifying effects on anthracycline dose-dependent cardiomyopathy risk (Wang et al., 2014; Wang et al., 2016).

Complex diseases are often characterized by many correlated traits which can better reflect their underlying mechanism. For example, hypertension can be characterized by systolic and diastolic blood pressure (Newton-Cheh et al. 2009); metabolic syndrome is evaluated by four component traits: high-density lipoprotein (HDL) cholesterol, plasma glucose and Type 2 diabetes, abdominal obesity, and diastolic blood pressure (Zabaneh and Balding, 2010); a person’s cognitive ability is usually measured by tests in domains including memory, intelligence, language, executive function, and visual-spatial function (Yang and Wang, 2012) and for chronic obstructive pulmonary disease (COPD), there are 7 key quantitative COPD-related phenotypes (Zhang et al., 2018). More and more large cohort studies have collected a broad array of correlated phenotypes to reveal the genetic components of many complex human diseases. Therefore, by jointly analyzing these correlated traits, we can not only gain more power by aggregating multiple weak effects, but also understand the genetic architecture of the disease of interest.

Evidence suggests that gene-environment interaction (GxE) effect may not only exist for a single trait but also exist for multiple traits of a disease. For example, a gene by stress genome-wide interaction analysis and path analysis identify EBF1 as a cardiovascular and metabolic risk gene. Gene EBF1 not only has gene-by-stress interaction effect for hip circumference but also has gene-by-stress interaction effects for waist circumference, body mass index (BMI), fasting glucose, type II diabetes status, and common carotid intimal medial thickness (CCIMT), supporting a model of gene-by-stress interaction that connects cardiovascular disease (CVD) risk factor endophenotypes such as central obesity and increased blood glucose or diabetes to CVD itself (Singh et al., 2015).

Examination of GxE for multiple traits can yield valuable insights about the etiology of the disease and increase power in detecting disease associated genes. Generally, a gene is the basic functional unit of inheritance. Thus, results from gene level association tests can be more readily integrated with downstream functional and pathogenic investigations. Next-generation sequencing technology provides potential opportunities to make directly testing both common and rare variants in a gene feasible (Andrés et al., 2007). Rare variants may play an important role in studying the etiology of complex human diseases where rare variants are usually defined as genetic variants with minor allele frequency (MAF) less than 5%. Numerous statistical methods have been developed for testing for rare variants, where gene-level analysis is often performed to jointly study the effects of rare variants in a gene such as the sequence kernel association test (SKAT) (Wu et al., 2011), the combined multivariate and collapsing (CMC) method (Li and Leal, 2008), the weighted sum statistic (WSS) (Madsen and Browning, 2009) and testing the effect of an Optimally Weighted combination of variants (TOW) (Sha et al., 2012).

The tests discussed above are designed to assess the association of the main effects of rare variants with a single trait. To our knowledge, limited methods have been developed for testing gene-environment interactions (GxEs) in sequencing association studies, especially for multiple traits. Existing methods for assessing common variants by environment interactions such as gene-environment set association test (GESAT) (Lin et al., 2013) will be less powerful when it is naively applied to rare variants gene-environment interaction analysis (Lin et al., 2016). To test rare variants by environment interactions, Lin et al. (2016) developed the interaction sequence kernel association test (ISKAT) to assess rare variants by environment interactions. When there is no prior information, ISKAT recommends Beta(MAF; 1, 25) as the weight which has the beta distribution density function with parameters 1 and 25 evaluated at the sample minor allele frequency (MAF). ISKAT may lose power when the MAFs of causal variants are not in the range (0.01,0.035).

Furthermore, existing methods for multiple-trait association tests have primarily focused on testing main effects of rare variants (Wu and James, 2016) or common variants (Ferreira and Purcell, 2008; Wu and Pankow, 2015). It is essential to develop novel methods for multiple traits to detect the gene-environment interactions for both rare and common variants.

In this article, we develop novel methods to test for rare and/or common variants by environment interactions on multiple traits in sequencing association studies. Our methods can be divided into three steps. We first use either principle component analysis or standardization analysis to release the correlation among the multiple traits. Then, we detect the effect of GxE for each transferred trait using novel proposed tests: testing the effect of an optimally weighted combination of GxE (TOW-GE) and/or variable weight TOW-GE (VW-TOW-GE). Both TOW-GE and VW-TOW-GE are robust to directions of effects of causal GxEs. Finally, we employ the Fisher’s combination test to combine the p-values of TOW-GE and/or VW-TOW-GE. Furthermore, we evaluate the performance of the proposed methods via simulation studies and real data analysis using the sequencing data from the COPDGene Study.

## Methods

Consider *n* unrelated individuals sequenced in a region with *m* variants and *K* measured continuous traits. For ease of presentation, we only consider a single environmental factor. We are interested in studying the *p* variants by environment interactions. The method can be easily extended to the case where there is more than one environmental factor. For individual *i* = 1, …, *n*, let ***Y***_*i*_ = (*y*_*i*1_, …, *y*_*iK*_)^*T*^ denote the *K* continuous traits, ***X***_*i*_ = (*x*_*i*1_, …, *x*_*iq*_)^*T*^ the q covariates, ***G***_*i*_ = (*g*_*i*1_, …, *g*_*ip*_)^*T*^ genotypes for the p rare variants in a genomic region (a gene or a pathway) and *E*_*i*_ environmental factor. For simplicity, we assume a common set of covariates for all traits which can be easily extended to the case of differing covariates. Let ***S***_*i*_ = (*E*_*i*_*g*_*i*1_, …, *E*_*i*_*g*_*ip*_)^*T*^ be a vector of variants by environment interaction terms, GxE, for the *i*^*th*^ individual.

The goal of our analysis is to study the association of these multiple phenotype traits with rare variants by environment interaction effects. Our method can be divided into the following three steps:

### Step 1

Denote the correlation matrix of ***Y***_*i*_ as **∑** = (*ρ*_*kl*_), where *ρ*_*kl*_ = *Cor*(*y*_*ik*_, *y*_*il*_) and define an *n* × *K* phenotype matrix ***Y*** = (***Y***_1_, …, ***Y***_*n*_)^*T*^. Due to the correlation among *y*_*i*1_, …, *y*_*iK*_, two methods can be used to reduce the correlation among them.

1. We propose to use principal component(PC) analysis for multiple traits and obtain independent phenotype matrix ***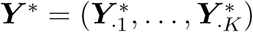***, where we denote ***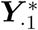*** as the first principal component and ***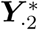*** as the second principle component and so on. According to the principal axis theorem in mechanics, we know that ***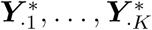*** are independent.
2. If we assume the mean and variance of ***Y***_*i*_ = (*y*_*i*1_, …, *y*_*iK*_)^*T*^ are ***µ*** and **∑**, then we can also obtain the independent vector ***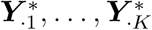*** by standardization: ***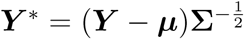***. In practice, we use the unbiased estimators of ***µ*** and **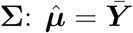** and **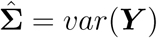**, where ***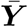*** denotes the sampe mean and *var*(***Y***) denotes the sample covariance matrix.

For simplicity of symbol, we will use *y*_*ik*_ to denote *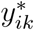*, the elements of ***Y***\*, in following paragraphs.

### Step 2

We use the generalized linear model to model the relationship between the *k*^*th*^ trait values *y*_*ik*_ and covariates ***X***_*i*_, genotypes ***G***_*i*_, environmental factor *E*_*i*_ and GxE ***S***_*i*_:

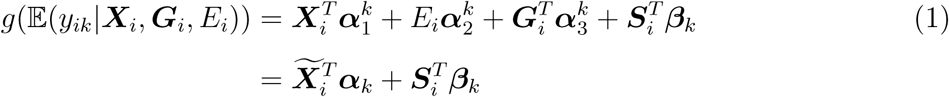

where *g*(·) be a canonical link function, ***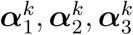*** and ***β***_*k*_ are defined as *q* × 1 coefficient vector of covariate, the coefficient of environmental factor, *p* × 1 coefficient vector of genotype and *p* × 1 coefficient vector of GxE interactions for the *i*^*th*^ individual and the *k*^*th*^ trait, respectively. Let ***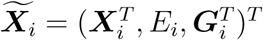*** and ***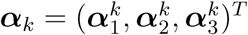***. For the *k*^*th*^ trait, we are interested in testing the null hypothesis *H*_0_ : ***β***_***k***_ = 0.

We develop a score test by treating ***α***_*k*_ as nuisance parameters and then adjust both the *k*^*th*^ trait value *y*_*ik*_ and ***S***_*i*_ for the covariates ***X***_*i*_, the genotypic score ***G***_*i*_, and the environmental variable *E*_*i*_ by applying linear regression. Denote *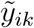* as the residual of *y*_*ik*_ and ***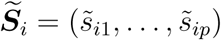*** as the residual of ***S***_*i*_. Then, the relationship between *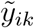* and ***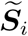*** can be modeled by the GLM:

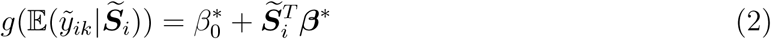

To test *H*_0_ : ***β***_***k***_ = 0 in (1) is equivalent to test *H*_0_ : ***β***\* = 0 in (2) (Sha et al. 2012). Here, we extend the TOW method proposed by Sha et al. (2012) to test the effect of a weighted combination of GxE, *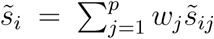*. Under the generalized linear model, the score test statistic given by Sha et al. (2011) becomes:

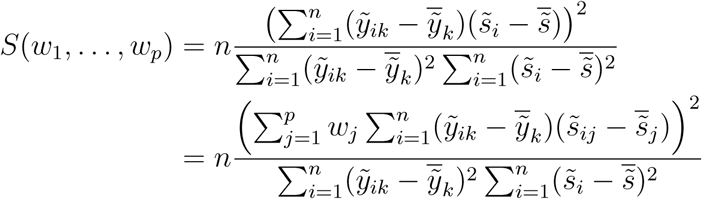

Because GxE for rare variants are essentially independent, we have:

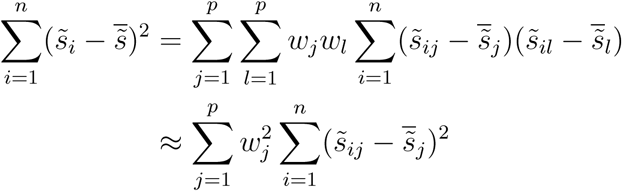

Thus, as a function of (*w*_1_, …, *w*_*p*_), the score test statistic *S*(*w*_1_, …, *w*_*p*_) reaches its maximum 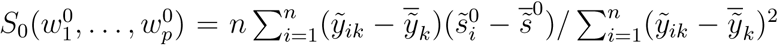 when *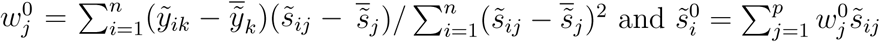*.

Similarly, we define the statistic to Test the effect of the Optimally Weighted combination of GxE (TOW-GE), 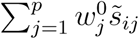 as:

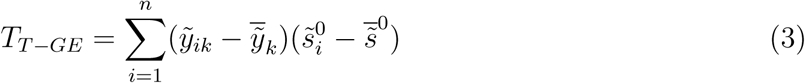

which is equivalent to *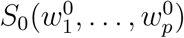* when we use a permutation test to evaluate p-values, where 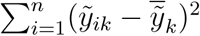 can be viewed as a constant.

The optimal weight *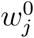* is equivalent to *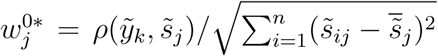* where *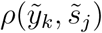* is the correlation coefficient between *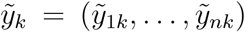* and *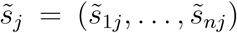*. From the expression of *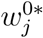*, we can see that it is proportional to *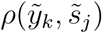* and thus *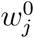* will put large weights to the GxE that have strong associations with the trait and also adjust for the direction of the association. Simultaneously, *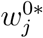* is proportional to 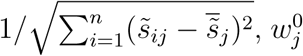 will put large weights to GxEs with small variations which are common in GxEs for rare variants.

TOW-GE focuses mainly on rare variants by environment interactions and it may lose power for both rare and common variants by environment interactions because it puts small weights on common variants by environment interactions. Thus, to test the GxE effects of both rare and common variants, we propose the following variable weight TOW-GE denoted as VW-TOW-GE. We first divide GxEs into two parts based on rare or common variants and then we apply TOW-GE to the two terms separately. Let 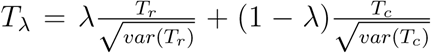 where *T*_*r*_ and *T*_*c*_ denote the test statistics of TOW-GE for GxE effects of rare and common variants, respectively. Denote *p*_*λ*_ as the p-value of *T*_*λ*_, and then the test statistic of VW-TOW-GE is defined as *T*_*VW -TOW -GE*_ = min_0*≤λ≤*1_ *p*_*λ*_. Here, we use a finite grid search of *λ* to evaluate the statistics *T*_*VW -TOW -GE*_. We divide the interval [0,1] into K subintervals of equal-length (*λ*_*k*_ = *k/K* for *k* = 0, …, *K*) and take the minimum p-value of *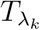* for a series of *λ*_*k*_. The p-value of *T*_*V W -T OW -GE*_ can be evaluated by permutation tests following similar steps of permutation tests for VW-TOW proposed by Sha et al. (2012).

### Step 3

For all K independent traits after transformed, we obtain K marginal p-values: *p*_1_, …, *p*_*K*_ from Step 2. Based on the Fisher’s combination test (FCT) (Fisher, 1932), the FCT for *K* traits is given by:

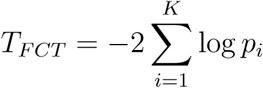

where *p*_*i*_ denotes the p-value of the test statistics *T*_*i*_ for the *i*th trait. Because these K traits are independent, the marginal p-values are independent and thus *T*_*F CT*_ follows a chi-squared distribution with 2*K* degrees of freedom.

## Comparison of tests

We compared the performance of our proposed methods with the interaction sequence kernel association test (ISKAT) (Lin et al., 2016), the modified WSS for testing GxEs (Madsen and Browning, 2009) and the modified CMC method for testing GxEs (Li and Leal, 2008) in Step 2. In this study, the rank sum test used by WSS and the *T*^2^ test used by CMC were replaced with the score test based on residuals *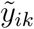* and *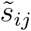*.

## Simulation

The empirical Mini-Exome genotype data provided by the GAW17 is used for simulation studies. The dataset contains genotypes of 697 unrelated individuals on 3,205 genes. Gene ELAVL4 in GAW17 was used to simulate GxE effect on quantitative trait *Q*_1_ which is follow normal distribution. *Q*_2_ is a quantitative trait which has weak correlation with *Q*_1_. We chose gene ELAVL4 in our simulation study. Gene ELAVL4 has 10 variants containing 8 rare variants and 2 common variants. Rare variants in the simulation are defined with MAF *<* 0.05.

We consider four correlated traits (K=4) with a compound-symmetry correlation matrix and consider two covariates: a standard normal covariate *X*_1_ and a binary covariate *X*_2_ with *P* (*X*_2_ = 1) = 0.5. The environmental factor E is assumed to be continuous following standard normal distribution. We generate trait values by using the following four models:

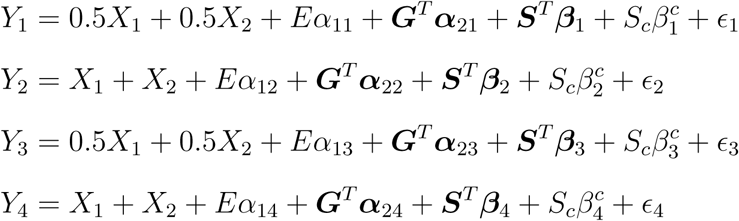

where *ϵ*_1_, *ϵ*_2_, *ϵ*_3_, *ϵ*_4_ are zero-mean normal with variances *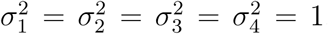*, correlation coefficient *ρ* = 0.5; ***α***_1_ = (0.015, 0.015, 0.03, *-*0.02); ***S*** is for the rare variants GxEs and *S*_*c*_ is one common variant GxE. Specifically, in the case of rare variants GxEs only, *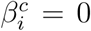* for *i* = 1, 2, 3, 4.

For type I error evaluation, we consider that there are no GxE effects in the aforementioned four models. Two scenarios are considered to evaluate the empirical type I error rates: (a) including main effects, we set the magnitudes of matrix ***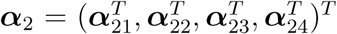*** as 0.3 and the sign for each ***α*** is randomly sampling from (*-*1, 1); (b) no main effects (***α***_2_ = 0).

For power comparisons, we consider that there are GxE effects in the aforementioned four models in two different cases: 1) rare variants GxEs only; 2) both rare and common variants GxE effects. We consider two different scenarios for each case: (a) main effects (***α***_2_ ≠ 0); (b) no main effects (***α***_2_ = 0). We vary the number of non-zero elements in vector ***β***_*i*_, proportion of non-zero elements in ***β***_*i*_ that are positive, and the magnitudes of the non-zero *β*_*ij*_. We set the magnitudes of the non-zero *β*_*ij*_’s as |*β*_*ij*_| = *c*, and increased c from 0.1 to 0.5.

## Simulation results

The empirical type I error rates are shown in Table 1 and Table 2. In each simulation scenario, p-values are estimated by 10,000 permutations and type I error rates are evaluated using 10,000 replicated samples. For 10,000 replicated samples, the 95% confidence intervals for type I error rates of nominal levels 0.05, 0.01 and 0.001 are (0.046, 0.054), (0.008, 0.012) and (0.0004, 0.0016), respectively. When there are (a) main effects (***α***_2_ ≠ 0), TOW-GE, VW-TOW-GE, ISKAT and WSS demonstrate well controlled type I error rates and the burden test CMC tends to have very conservative type I error rates (top panel of Table 1 and Table 2). When there are (b) no main effects (***α***_2_ = 0), all methods control type I error rates very well (bottom panel of Table 1 and Table 2).

**Table 1:** Type 1 error rates for rare variants in the presence of main effects (top panel) and in the absence of main effects (bottom panel) for n=2000, where the type I error out of parentheses based on PC analysis and the type I error in parentheses based on standardization in Step 1.

| With main effect |  |  |  |  |  |
| --- | --- | --- | --- | --- | --- |
| | α-level | TOW-GE | ISKAT | WSS | CMC |
| $n = 2000$ | 0.05 | 0.059(0.063) | 0.062(0.058) | 0.057(0.054) | 0.021(0.023) |
|  | 0.01 | 0.013(0.015) | 0.017(0.020) | 0.015(0.018) | 0.001(0.002) |
|  | 0.001 | 0.004(0.003) | 0.007(0.009) | 0.002(0.004) | 0.000(0.000) |
| Without main effect |  |  |  |  |  |
| $n = 2000$ | 0.05 | 0.051(0.050) | 0.060(0.062) | 0.053(0.062) | 0.042(0.046) |
|  | 0.01 | 0.011(0.013) | 0.016(0.014) | 0.009(0.013) | 0.009(0.008) |
|  | 0.001 | 0.003(0.002) | 0.006(0.006) | 0.001(0.002) | 0.002(0.001) |

**Table 2:**
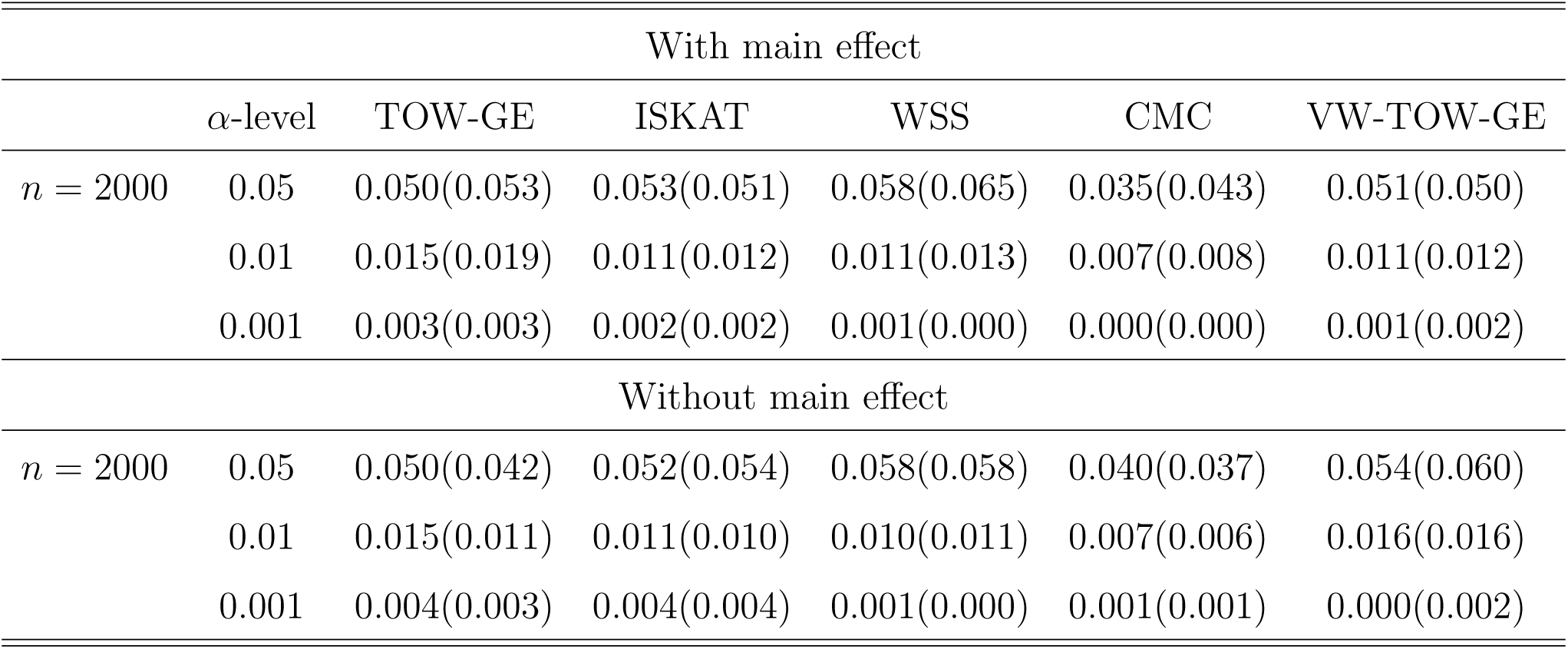
Type 1 error rates for both rare and common variants in the presence of main effects (top panel) and in the absence of main effects (bottom panel) for n=2000, where the type I error out of parentheses based on PC analysis and the type I error in parentheses based on standardization according to Step 1.

| With main effect |  |  |  |  |  |  |
| --- | --- | --- | --- | --- | --- | --- |
| | α-level | TOW-GE | ISKAT | WSS | CMC | VW-TOW-GE |
| $n = 2000$ | 0.05 | 0.050(0.053) | 0.053(0.051) | 0.058(0.065) | 0.035(0.043) | 0.051(0.050) |
|  | 0.01 | 0.015(0.019) | 0.011(0.012) | 0.011(0.013) | 0.007(0.008) | 0.011(0.012) |
|  | 0.001 | 0.003(0.003) | 0.002(0.002) | 0.001(0.000) | 0.000(0.000) | 0.001(0.002) |
| Without main effect |  |  |  |  |  |  |
| $n = 2000$ | 0.05 | 0.050(0.042) | 0.052(0.054) | 0.058(0.058) | 0.040(0.037) | 0.054(0.060) |
|  | 0.01 | 0.015(0.011) | 0.011(0.010) | 0.010(0.011) | 0.007(0.006) | 0.016(0.016) |
|  | 0.001 | 0.004(0.003) | 0.004(0.004) | 0.001(0.000) | 0.001(0.001) | 0.000(0.002) |

Power comparisons of four tests (TOW-GE, ISKAT, WSS and CMC) for sample size n from 1000 to 4000 at significance level *α* = 0.05 for testing rare variants GxE effects on continuous traits with main effect are shown in Table 3 and without main effect are shown in Table 4. From these two tables, we can see that the empirical power for all methods are increasing when the sample size *n* increases. When the sample size *n* = 2000, we can easily see the difference regarding the power of different methods.

**Table 3:** Power comparisons of four tests (TOW-GE, ISKAT, WSS and CMC) for different samples n at *α* = 0.05 level of significance for testing rare variant by environment interaction effects on a continuous outcome with main effect where we set 50% of the *β*_*ij*_ are positive and the power out of parentheses based on PC analysis and the power in parentheses based on standardization.

| With main effect |  |  |  |  |  |
| --- | --- | --- | --- | --- | --- |
| | $\vert \beta\vert $ | TOW-GE | ISKAT | WSS | CMC |
| n=1000 | 0.1 | 0.091(0.086) | 0.085(0.073) | 0.087(0.080) | 0.024(0.026) |
|  | 0.2 | 0.153(0.117) | 0.148(0.122) | 0.148(0.107) | 0.066(0.057) |
|  | 0.3 | 0.348(0.235) | 0.278(0.221) | 0.314(0.233) | 0.135(0.079) |
|  | 0.4 | 0.574(0.399) | 0.444(0.348) | 0.464(0.346) | 0.253(0.159) |
|  | 0.5 | 0.802(0.611) | 0.543(0.470) | 0.623(0.490) | 0.357(0.235) |
| n=2000 | 0.1 | 0.091(0.080) | 0.109(0.087) | 0.093(0.079) | 0.050(0.041) |
|  | 0.2 | 0.292(0.203) | 0.243(0.179) | 0.181(0.131) | 0.113(0.069) |
|  | 0.3 | 0.589(0.384) | 0.471(0.378) | 0.355(0.238) | 0.261(0.171) |
|  | 0.4 | 0.903(0.729) | 0.555(0.493) | 0.538(0.420) | 0.459(0.298) |
|  | 0.5 | 0.986(0.916) | 0.627(0.587) | 0.673(0.561) | 0.646(0.478) |
| n=3000 | 0.1 | 0.123(0.097) | 0.129(0.107) | 0.092(0.082) | 0.062(0.049) |
|  | 0.2 | 0.385(0.240) | 0.357(0.267) | 0.179(0.131) | 0.145(0.088) |
|  | 0.3 | 0.808(0.601) | 0.542(0.474) | 0.372(0.274) | 0.377(0.232) |
|  | 0.4 | 0.982(0.908) | 0.640(0.597) | 0.582(0.452) | 0.648(0.485) |
|  | 0.5 | 0.999(0.990) | 0.695(0.644) | 0.717(0.604) | 0.841(0.701) |
| n=4000 | 0.1 | 0.141(0.108) | 0.154(0.117) | 0.093(0.085) | 0.062(0.043) |
|  | 0.2 | 0.536(0.339) | 0.448(0.354) | 0.204(0.148) | 0.203(0.137) |
|  | 0.3 | 0.936(0.786) | 0.630(0.580) | 0.406(0.286) | 0.468(0.310) |
|  | 0.4 | 0.999(0.981) | 0.639(0.588) | 0.591(0.457) | 0.790(0.634) |
|  | 0.5 | 1.000(0.999) | 0.750(0.692) | 0.767(0.659) | 0.916(0.815) |

**Table 4:** Power comparisons of four tests (TOW-GE, ISKAT, WSS and CMC) for different samples n at *α* = 0.05 level of significance for testing rare variant by environment interaction effects on a continuous outcome with no main effect where we set 50% of the *β*_*ij*_ are positive and the power out of parentheses based on PC analysis and the power in parentheses based on standardization.

| With no main effect |  |  |  |  |  |
| --- | --- | --- | --- | --- | --- |
| | $\vert \beta\vert $ | TOW-GE | ISKAT | WSS | CMC |
| n=1000 | 0.1 | 0.076(0.083) | 0.099(0.092) | 0.063(0.058) | 0.049(0.043) |
|  | 0.2 | 0.127(0.099) | 0.162(0.117) | 0.102(0.092) | 0.081(0.076) |
|  | 0.3 | 0.323(0.209) | 0.307(0.231) | 0.203(0.134) | 0.174(0.114) |
|  | 0.4 | 0.573(0.394) | 0.466(0.366) | 0.309(0.227) | 0.292(0.210) |
|  | 0.5 | 0.810(0.617) | 0.528(0.460) | 0.429(0.333) | 0.383(0.272) |
| n=2000 | 0.1 | 0.084(0.073) | 0.095(0.079) | 0.075(0.071) | 0.074(0.069) |
|  | 0.2 | 0.283(0.191) | 0.254(0.191) | 0.150(0.111) | 0.164(0.104) |
|  | 0.3 | 0.588(0.389) | 0.453(0.346) | 0.239(0.166) | 0.290(0.214) |
|  | 0.4 | 0.912(0.730) | 0.581(0.519) | 0.394(0.295) | 0.461(0.344) |
|  | 0.5 | 0.991(0.921) | 0.659(0.612) | 0.532(0.405) | 0.606(0.458) |
| n=3000 | 0.1 | 0.117(0.092) | 0.127(0.098) | 0.081(0.083) | 0.092(0.070) |
|  | 0.2 | 0.391(0.236) | 0.375(0.277) | 0.149(0.111) | 0.198(0.136) |
|  | 0.3 | 0.819(0.605) | 0.546(0.467) | 0.296(0.211) | 0.403(0.283) |
|  | 0.4 | 0.986(0.915) | 0.646(0.613) | 0.479(0.354) | 0.629(0.489) |
|  | 0.5 | 1.000(0.996) | 0.720(0.657) | 0.612(0.497) | 0.801(0.693) |
| n=4000 | 0.1 | 0.142(0.113) | 0.153(0.110) | 0.087(0.074) | 0.092(0.079) |
|  | 0.2 | 0.546(0.344) | 0.425(0.327) | 0.167(0.129) | 0.245(0.180) |
|  | 0.3 | 0.952(0.803) | 0.632(0.571) | 0.344(0.237) | 0.502(0.369) |
|  | 0.4 | 0.999(0.980) | 0.689(0.622) | 0.501(0.387) | 0.765(0.632) |
|  | 0.5 | 1.000(1.000) | 0.792(0.718) | 0.668(0.541) | 0.878(0.784) |

The results for *n* = 2000 are also given in Figure 1 and Figure 2 including main effect and no main effect for rare variants GxE effects based on PC analysis and Figure 3 and Figure 4 including main effect and no main effect for rare variants GxE effects based on standardization, respectively. The top, middle, and bottom panels of these figures correspond to the three scenarios in which there are 2, 6 and 8 non-zero *β*_*ij*_’s, respectively. The left and right panels of figures correspond to two cases in which 50% of the *β*_*ij*_ are positive and 100% of the *β*_*ij*_ are positive for the *i*th trait, respectively. For each plot, we vary c, the magnitudes of the non-zero *β*_*ij*_. As shown in the four figures for the case where 50% of the *β*_*ij*_ are positive, TOW-GE is more powerful than the other three tests. For the case where 100% of the *β*_*ij*_ are positive, WSS is relatively more powerful than TOW-GE and TOW-GE is more powerful than the other two tests. The WSS is very sensitive to the directions of effects due to its simple aggregation of GxEs. Among the four tests (TOW-GE, ISKAT, WSS and CMC) in the two different cases, CMC is the least powerful test.

**Figure 1.**
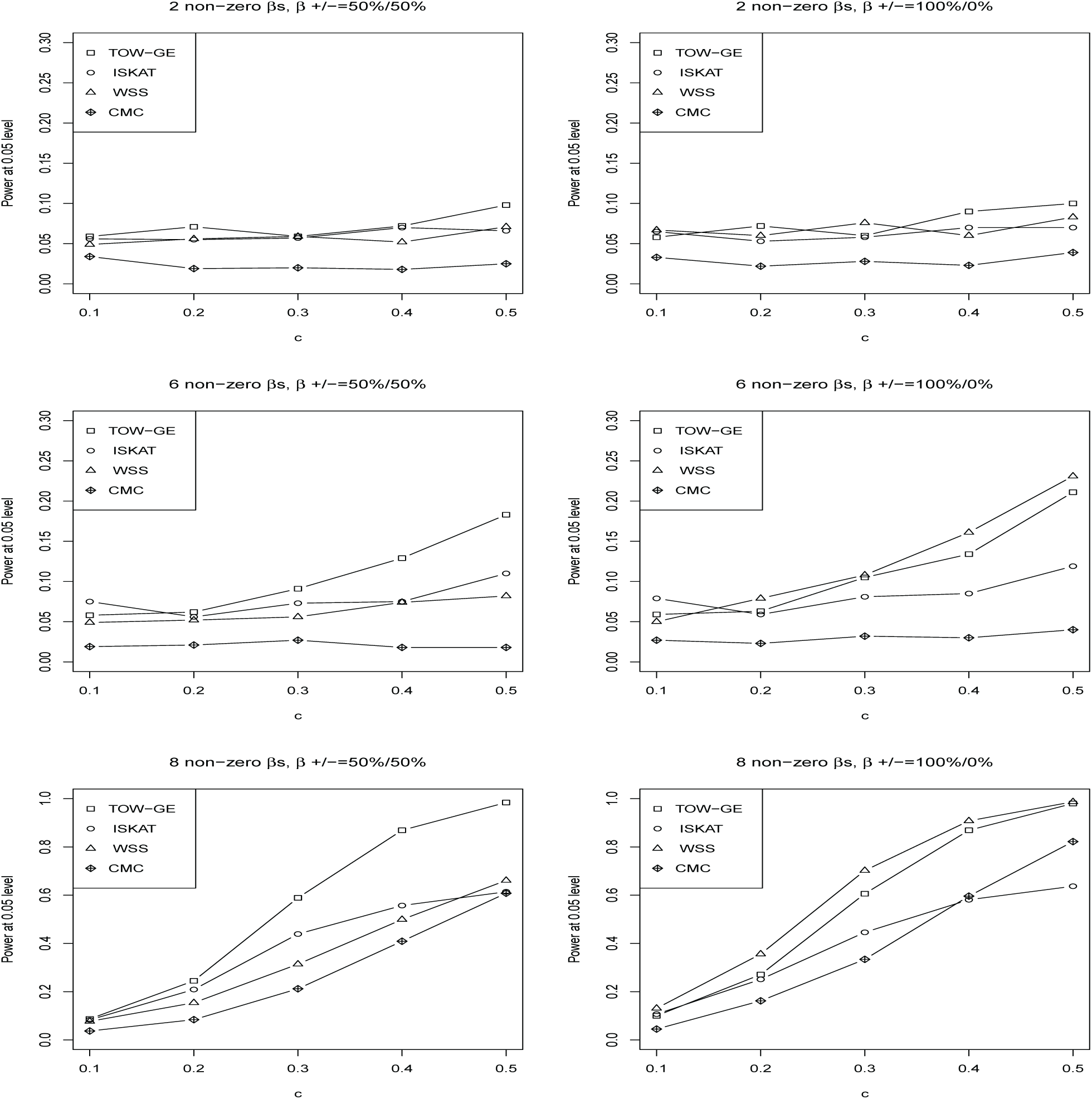
Power comparisons of four tests (TOW-GE, ISKAT, WSS and CMC) for n=2000 at *α* = 0.05 level of significance for testing rare variant by environment interaction effects on a continuous outcome when there are main effects based on PC analysis.

**Figure 2.**
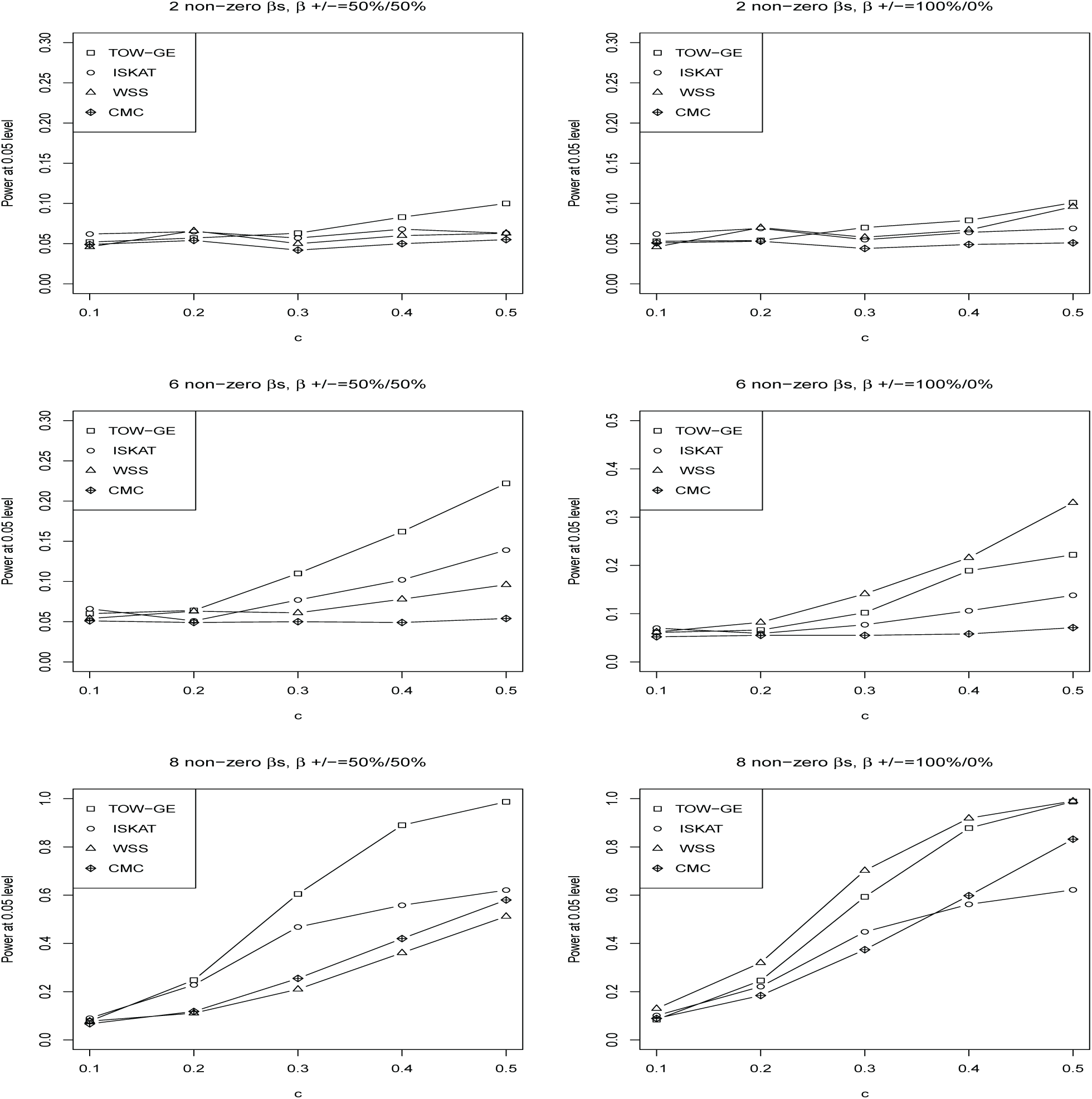
Power comparisons of four tests (TOW-GE, ISKAT, WSS and CMC) for n=2000 at *α* = 0.05 level of significance for testing rare variant by environment interaction effects on a continuous outcome when there are no main effects based on PC analysis.

**Figure 3.**
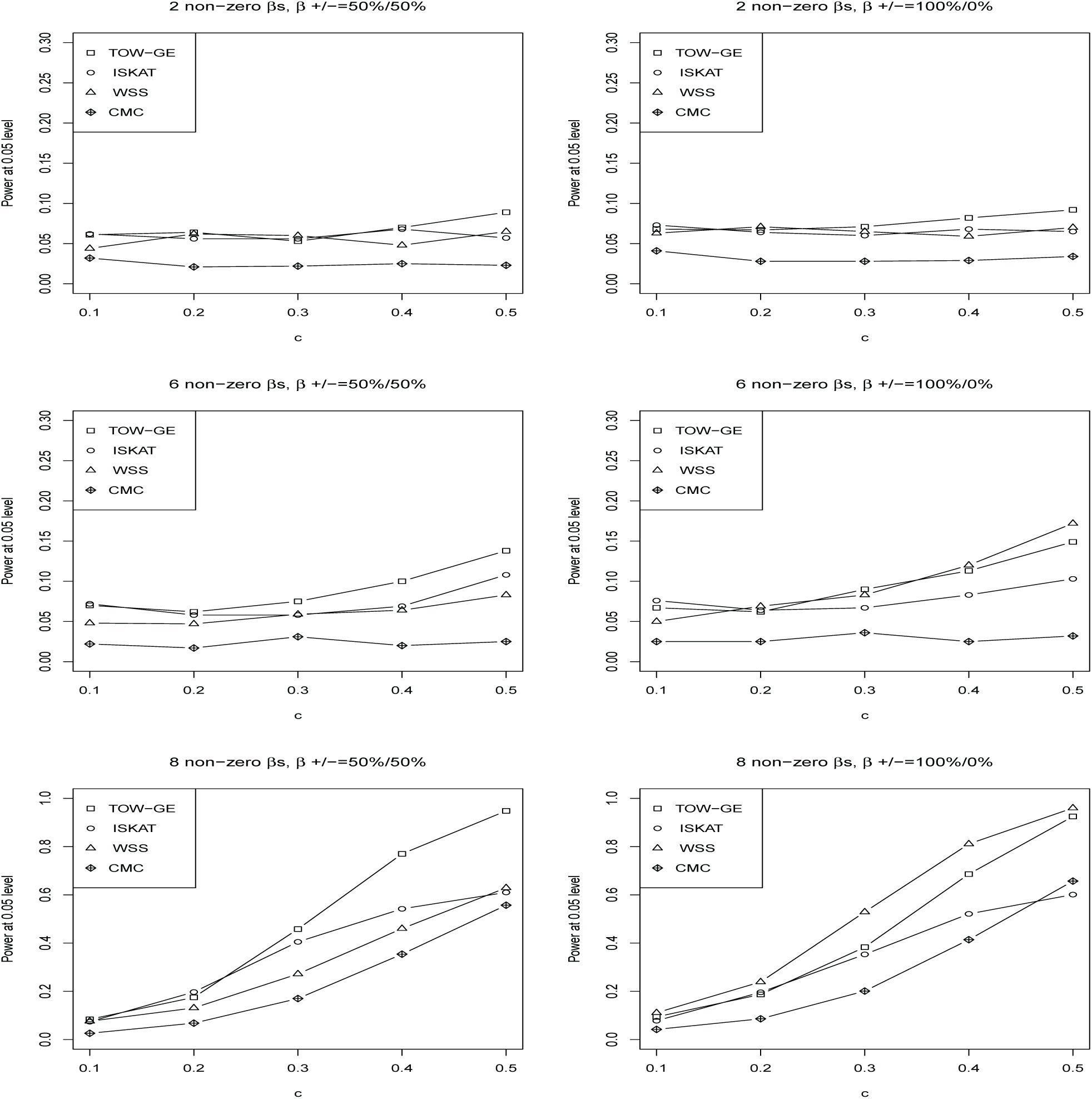
Power comparisons of four tests (TOW-GE, ISKAT, WSS and CMC) for n=2000 at *α* = 0.05 level of significance for testing rare variant by environment interaction effects on a continuous outcome when there are main effects based on standardization.

**Figure 4.**
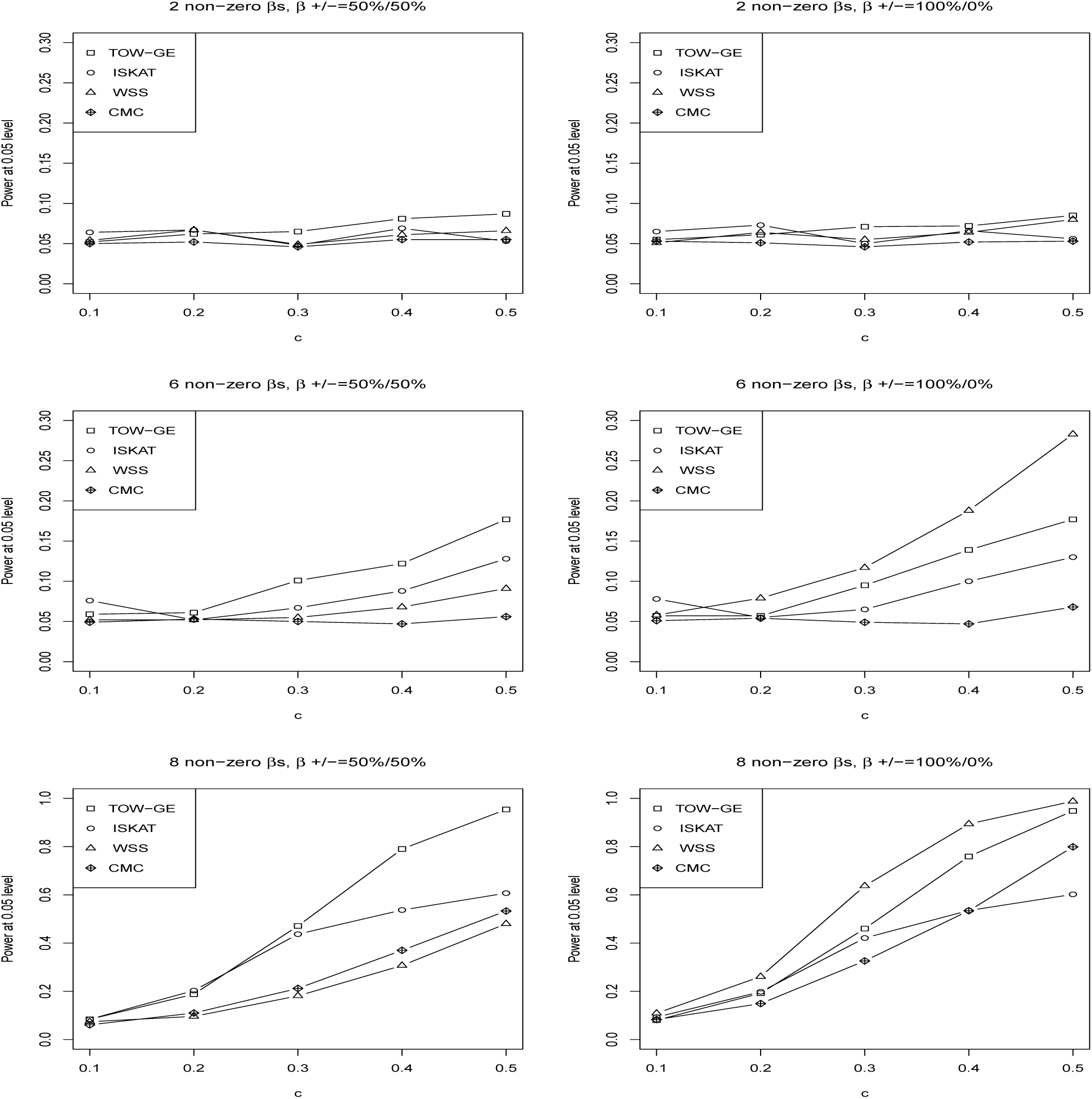
Power comparisons of four tests (TOW-GE, ISKAT, WSS and CMC) for n=2000 at *α* = 0.05 level of significance for testing rare variant by environment interaction effects on a continuous outcome when there are no main effects based on standardization.

Power comparisons of the five tests (TOW-GE, VW-TOW-GE, ISKAT, WSS and CMC) for both rare and common variants GxE effects are given in Figure 5. For each plot, we vary c from 0.02 to 0.1 and set 50% of the *β*_*ij*_ as positive. Simultaneously, we set the coefficient of common variant by interaction *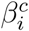* as positive and the magnitude of *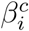* as twice of *β*_*ij*_. From Figure 5, we can see that the empirical power of TOW-GE, VW-TOW-GE and CMC outperform the other two tests and WSS is the least powerful test. WSS loses power because it gives common variant by environment interaction effect very small weight. The method ISKAT performs worse than our methods, it is similar to SKAT, which will lose power when the MAFs of causal variants are not in the range (0.01,0.035) (Yang et al. 2017).

**Figure 5.**
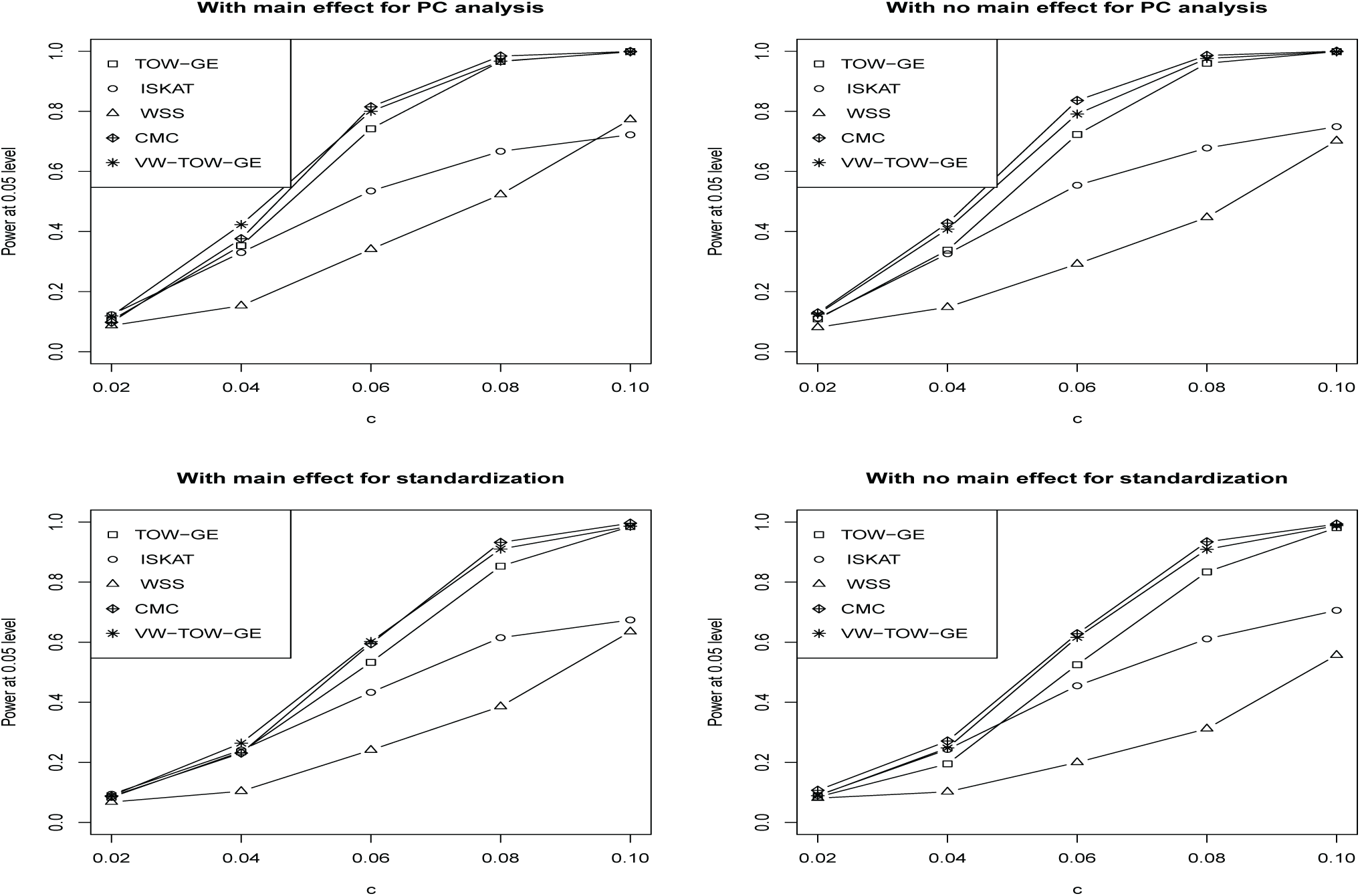
Power comparisons of five tests (TOW-GE, ISKAT, WSS,CMC and VW-TOW-GE) for n=2000 at *α* = 0.05 level of significance for testing both rare and common variant by environment interaction effects on a continuous outcome. Top panel: PC analysis; bottom panel: Standardization; Left panel: With main effect; Right panel: With no main effect.

In order to understand when and which approach should be recommended, power comparisons between PC analysis (PCA) and Standardization (STD) analysis for TOW-GE are performed. Power of TOW-GE for n=2000 at *α* = 0.05 level of significance for testing rare variant environment interaction effects on a continuous trait with main effect and without main effect are shown in Figure 6 and Figure 7, respecively. As shown in these figures, we can see that TOW-GE using PCA is relatively more powerful than TOW-GE using STD.

**Figure 6.**
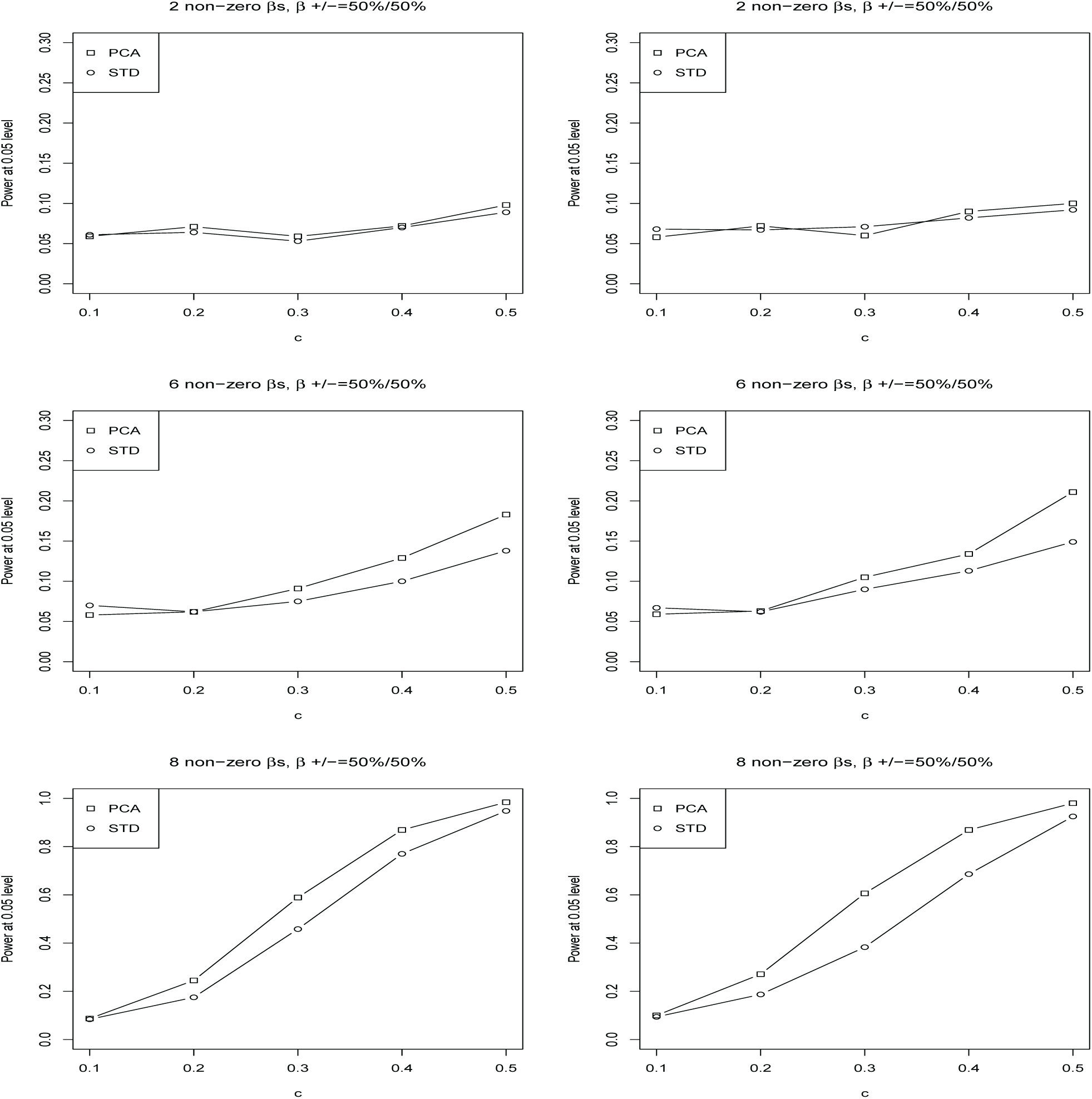
Power comparisons for PCA and Standardization(STD) of test TOW-GE for n=2000 at *α* = 0.05 level of significance for testing rare variant by environment interaction effects on a continuous outcome with main effect.

**Figure 7.**
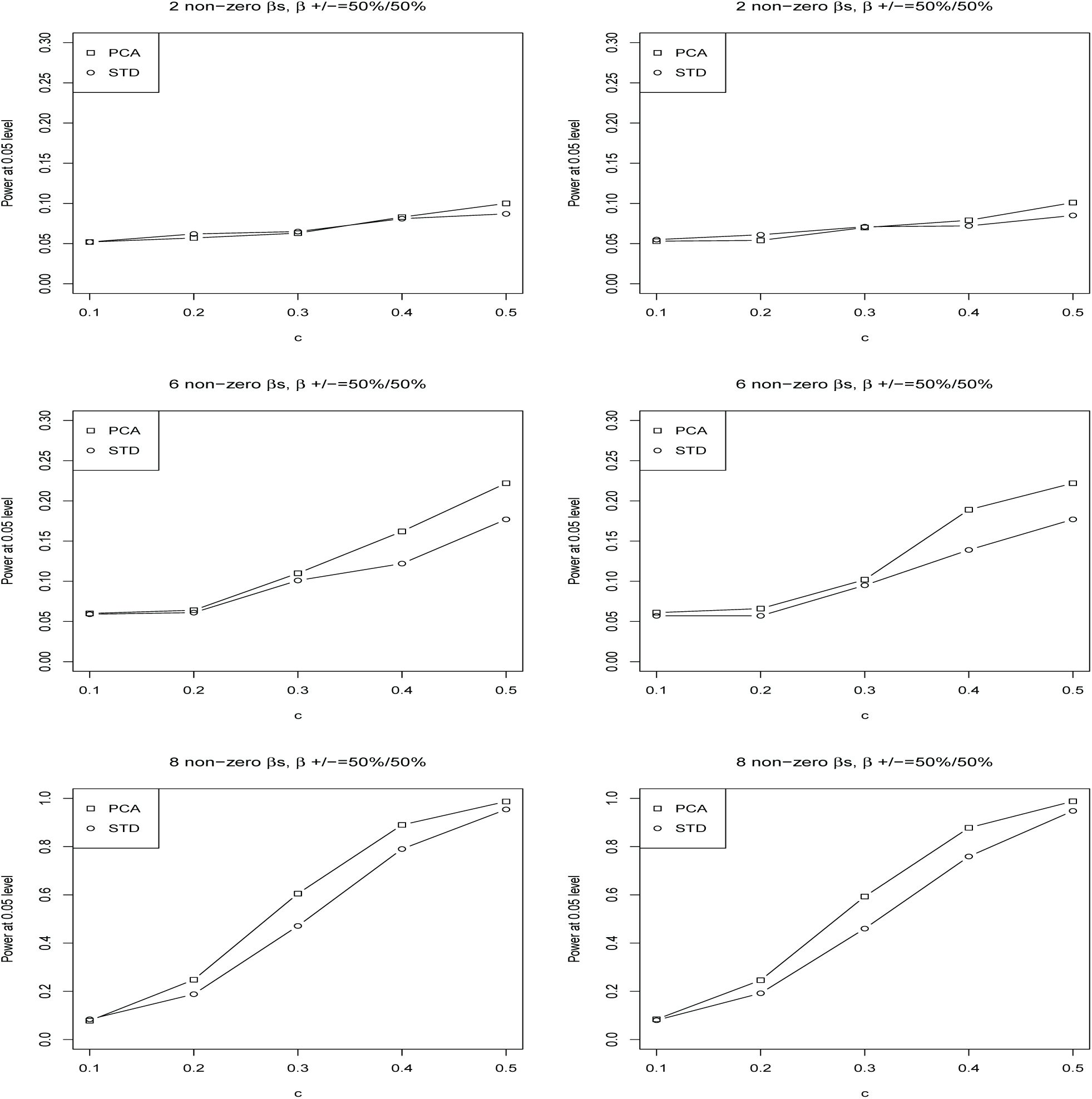
Power comparisons for PCA and Standardization(STD) of test TOW-GE for n=2000 at *α* = 0.05 level of significance for testing rare variant by environment interaction effects on a continuous outcome with no main effect.

In summary, TOW-GE is the most powerful test in the case of rare variants by environment interaction effects except for the case where 100% of the *β*_*ij*_ are positive. VW-TOW-GE is either the most powerful test or has similar power to the most powerful test in the case of both rare and common variants GxE effects.

## Real data analysis

Chronic obstructive pulmonary disease (COPD) is one of the most common lung diseases characterized by long term poor airflow. COPD affects millions of Americans and is the third leading cause of disease-related death in the U.S. (Murphy and Sethi 2002). It is a complex disease influenced by genetic factors, environmental factors, and gene-environment interactions. It is well known that cigarette smoking is the main causal factor of COPD (Sandford and Silverman, 2002). Several genes are suggested to play a role in the presence of a gene-by-smoking interaction. For example, Hersh et al. (2005) reported that the SFTPB Thr131Ile polymorphism was associated with COPD, but only in the presence of a gene by environment interaction term, the SNP rs2292566 in gene EPHX1 was associated with COPD only in presence of a gene-by-smoking (pack-years) interaction term, and the 30-repeat allele of HMOX1 was associated. Celedon et al. (2004) showed that two SNPs in the promoter region of TGFB1 (rs2241712 and rs1800469) and one SNP in exon 1 of TGFB1 (rs1982073) were significantly associated with COPD among smokers in a COPD case control study. Sandford and Silverman (2002) presented that the GSTM1 gene was associated with severe chronic bronchitis in heavy smokers and an association of the TNF -308A allele with COPD was found in a Taiwanese population.

The COPDGene Study is a multi-center genetic and epidemiologic investigation to study COPD (Regan et al., 2011). This study is sufficiently large and appropriately designed for analysis of COPD. In this study, we consider more than 5000 non-Hispanic Whites (NHW) participants where the participants have completed a detailed protocol, including questionnaires, pre- and post-bronchodilator spirometry, high-resolution CT scanning of the chest, exercise capacity (assessed by six-minute walk distance), and blood samples for genotyping. The participants were genotyped using the Illumina OmniExpress platform. The genotype data has gone through standard quality-control procedures for genome-wide association analysis detailed at http://www.copdgene.org/sites/default/files/GWAS_QC_Methodology_20121115.pdf. We performed genotype imputation using 1000 Genomes Phase I v3 European reference panel.

Based on the literature studies of COPD (Chu et al., 2014 and Han et al., 2011), we selected 7 key quantitative COPD-related phenotypes, including FEV1 (% predicted FEV1), Emphysema (Emph), Emphysema Distribution (EmphDist), Gas Trapping (GasTrap), Airway Wall Area (Pi10), Exacerbation frequency (ExacerFreq), Six-minute walk distance (6MWD), 3 covariates, including BMI, age, and sex, and one environmental factor (Pack-Years). EmphDist is the ratio of emphysema at -950 HU in the upper 1*/*3 of lung fields compared to the lower 1*/*3 of lung fields where we did a log transformation on EmphDist in the following analysis, referred to Chu et al. (2014). In the analysis, participants with missing data in any of these phenotypes were excluded.

To evaluate the performance of our proposed method on a real data set, we applied all of the 5 methods (TOW-GE, ISKAT, WSS, CMC and VW-TOW-GE) to six COPD associated genes (SFTPB, EPHX1, GSTM1, HMOX1, TGFB1 and TNF) through an interaction with cigarette smoking. In the analysis, we excluded the extreme low frequency SNPs (MAF*<*0.001) and individuals with missing values in any of the 7 phenotypes and 3 covariates. We considered three different scenarios: (1) gene main effects, (2) gene-by-smoking interaction with main effect, and (3) gene-by-smoking interaction without main effect. When we considered only main effects of genes, we used five popular genetic association testing methods (TOW, SKAT, WSS, CMC and VW-TOW) in Step 2. We adopted 10^4^ permutations for our methods and used 0.05 as the significance level. The results are summarized in Table 5. From Table 5, we can see that two genes (SFTPB and GSTM1) are only identified by our methods and the gene HMOX1 is identified by SKAT and WSS methods for scenario (1) gene main effects; gene TNF is only identified by CMC method for scenario (2) gene-by-smoking interaction with main effect; three genes (SFTPB, HMOX1 and TGFB1) are identified by our methods and the HMOX1 gene is also identified by ISKAT and WSS methods for scenario (3), gene-by-smoking interaction without main effect. Our method identified most of the COPD associated genes especially in scenario (3).

**Table 5:** Summary of association analysis results of the COPD dataset. The p-value is calculated for gene with main effects (top panel), gene-by-smoking interaction with main effect (middle panel), Gene-by-smoking interaction without main effect (bottom panel) where p-value out of parentheses based on PC analysis and p-value in parentheses based on standardization according to Step 1.

| Gene main effects |  |  |  |  |  |
| --- | --- | --- | --- | --- | --- |
| Gene name | TOW | SKAT | WSS | CMC | VW-TOW |
| SFTPb | **0.0421(0.0456)** | 0.3321(0.3312) | 0.2008(0.3051) | 0.7087(0.8450) | 0.0891(0.2553) |
| EPHX1 | 0.5026(0.5070) | 0.3755(0.3385) | 0.9814(0.9774) | 0.8803(0.9100) | 0.5902(0.3886) |
| GSTM1 | **0.0235(0.0252)** | 0.3234(0.3151) | 0.6725(0.7374) | 0.3630(0.3038) | **0.0263(0.0586)** |
| HMOX1 | 0.1024(0.0858) | **0.0315(0.0596)** | **0.0067(0.0203)** | 0.9999(0.9997) | 0.3001(0.1994) |
| TGFB1 | 0.0825(0.0798) | 0.1128(0.1695) | 0.4454(0.4591) | 0.2170(0.3686) | 0.4556(0.1576) |
| TNF | 0.2154(0.2074) | 0.2198(0.2346) | 0.0879(0.2169) | 0.7663(0.7416) | 0.3094(0.2562) |
| Gene-by-smoking interaction with main effect |  |  |  |  |  |
| Gene name | TOW-GE | ISKAT | WSS | CMC | VW-TOW-GE |
| SFTPb | 0.3392(0.2733) | 0.6413(0.6035) | 0.6253(0.5863) | 0.8587(0.8006) | 0.4688(0.6582) |
| EPHX1 | 0.5742( 0.5757) | 0.5533(0.5174) | 0.3550(0.3527) | 0.9870(0.9722) | 0.8870(0.9833) |
| GSTM1 | 0.5735(0.5074) | 0.2983(0.2542) | 0.9480(0.9074) | 0.3397(0.3765) | 0.5520(0.5297) |
| HMOX1 | 0.4390( 0.4341) | 0.2395(0.2196) | 0.2360(0.1408) | 0.9999(0.9997) | 0.5692(0.4876) |
| TGFB1 | 0.4020(0.4195) | 0.6064(0.5688) | 0.4755(0.5102) | 0.9903(0.9981) | 0.3891(0.4833) |
| TNF | 0.8295(0.8619) | 0.6049(0.6282) | 0.1669(0.2875) | **0.0305(0.0427)** | 0.8572(0.8884) |
| Gene-by-smoking interaction without main effect |  |  |  |  |  |
| Gene name | TOW-GE | ISKAT | WSS | CMC | VW-TOW-GE |
| SFTPb | **0.0026(0.0012)** | 0.1048(0.0913) | 0.1089(0.0908) | 0.3569(0.4727) | **0.0214(0.0270)** |
| EPHX1 | 0.2298(0.2100) | 0.2307(0.2047) | 0.9178(0.9264) | 0.4418(0.4286) | 0.3199(0.3834) |
| GSTM1 | 0.0825(0.0867) | 0.1741(0.1983) | 0.6195(0.7102) | 0.3092(0.3279) | 0.0898(0.1780) |
| HMOX1 | **0.0474(0.0481)** | **0.0177(0.0099)** | **0.0009(0.0013)** | 0.9994(0.9985) | 0.0839(0.0982) |
| TGFB1 | **0.0290(0.0374)** | 0.0786(0.0913) | 0.4781(0.5078) | 0.1920(0.2101) | 0.1130(0.0705) |
| TNF | 0.2396(0.2372) | 0.2218(0.2470) | 0.0949(0.1793) | 0.2010(0.2047) | 0.4084(0.3226) |

Note: The bold numbers represent significant p-value for significance level 0.05.

## Discussion

Complex diseases are often characterized by correlated traits. For an example, hypertension, is defined by two levels according to the American College of Cardiology/American Heart Association (ACC/AHA) guidelines: (1) elevated blood pressure, with a systolic pressure (SBP) between 120 and 129 mm Hg and diastolic pressure (DBP) less than 80 mm Hg, and (2) stage 1 hypertension, with an SBP of 130 to 139 mm Hg or a DBP of 80 to 89 mm Hg (Whelton et al., 2018). Combining multiple correlated traits in genetic association test can not only enhance our understanding of the etiology of the disease but also increase power in testing GxE interaction effects. In this paper, we propose a new framework to test the association between GxEs and multiple traits. The proposed method can be divided into three steps: 1) reduce correlation among traits; 2) test GxE effect for each transferred trait; 3) combine p-values of all transfered traits by Fisher’s combination test. PC analysis and Standardization analysis are applied to reduce the correlation among the correlated traits in Step 1. Then, we apply extended TOW method to test rare variants by environment interaction (TOW-GE) and VW-TOW method to test both rare and common variants by environment interaction (VW-TOW-GE) in Step 2. We use the simulation studies to demonstrate that the proposed methods outperform the comparable methods in most of the scenarios with or without main effect. In addition, we test GxE effects for 7 related phenotypes of COPD. Our proposed methods verified the largest number of genes especially for gene-by-smoking interactions without main effect.

We noticed that the CMC loses power especially when the regions tested are large because of the increased degrees of freedom for Hotellings T2 such as gene HMOX1 in COPD data (13,854 bases).The CMC is powerful when directions of GxE effects of all causal variants are the same and the gene tested is short such as gene TNF (2,770 bases).

In this paper, we focus on the joint analysis of multiple continuous traits with GxE effects. Our results show that the proposed methods TOW-GE or VW-TOW-GE are the most powerful tests or have similar power to the most powerful test compared with competing methods. It will be interesting to generalize the methods to study the joint test of mixed type of phenotypes, which is analytically and computationally more challenging. TOW-GE is similar to TOW, which is derived for independent variants. Since common variants within a gene are usually correlated, we wonder the performance of TOW-GE for common variant GxE interaction effects. It will be of future research interest to assess the performance of TOW-GE for common variant GxE interaction effects.

## Acknowledgements

Q. Sha was supported by the National Human Genome Research Institute of the National Institutes of Health under Award Number R15HG008209. X. Raymond Gao was supported by National Institutes of Health (NIH; Bethesda, MD, USA) grants R01EY027315 and R-F1AG060472. The content is solely the responsibility of the authors and does not necessarily represent the official views of the National Institutes of Health. X. Wang was supported by the University of North Texas Foundation which was contributed by Dr. Linda Truitt Creagh. The content is solely the responsibility of the authors and does not necessarily represent the views of the University of North Texas Foundation and Dr. Linda Truitt Creagh. The Genetic Analysis workshops are supported by NIH grant R01 GM031575 from the National Institute of General Medical Sciences. Preparation of the Genetic Analysis Workshop 17 Simulated Exome Data Set was supported in part by NIH R01 MH059490 and used sequencing data from the 1000 Genomes Project (www.1000genomes.org). This research used data generated by the COPDGene study (phs000179/HMB and phs000179/DS-CS-RD), which was supported by National Institutes of Health (NIH) grants U01HL089856 and U01HL089897. The content is solely the responsibility of the authors and does not necessarily represent the official views of the National Heart, Lung, and Blood Institute or the National Institutes of Health. The COPDGene project is also supported by the COPD Foundation through contributions made by an Industry Advisory Board comprised of Pfizer, AstraZeneca, Boehringer Ingelheim, Novartis, and Sunovion.

